# Temperature as a likely driver shaping global patterns in mineralogical composition in bryozoans: Implications for marine calcifiers under Global Change

**DOI:** 10.1101/2022.09.30.510275

**Authors:** Blanca Figuerola, Huw J. Griffiths, Malgorzata Krzeminska, Anna Piwoni-Piorewicz, Anna Iglikowska, Piotr Kuklinski

## Abstract

The Southern Ocean is showing one of the most rapid responses to human-induced global change, thus acting as a sentinel of the effects on marine species and ecosystems. Ocean warming and acidification are already impacting benthic species with carbonate skeletons, but the magnitude of these changes to species and ecosystems remains largely unknown. Here we provide the largest carbonate mineralogical dataset to date for Southern Ocean bryozoans, which are diverse, abundant and important as carbonate producers, thus making them excellent for monitoring the effects of ocean warming and acidification. To improve our understanding of how bryozoans might respond to ocean warming and acidification, we assess latitudinal and seafloor temperature patterns of skeletal mineralogy using bryozoan species occurrences together with temperature data for the first time. Our findings, combining new mineralogical data with published data from warmer regions, show that the proportions of high-Mg calcite and bimineralic species increase significantly towards lower latitudes and with increasing seawater temperature. These patterns are consistent with the hypothesis that seawater temperature is likely a significant driver of variations in bryozoan mineralogy at a global scale.

## 1 INTRODUCTION

Increased levels of atmospheric carbon dioxide (CO_2_) as a result of human activities are driving rising temperatures, ocean acidification and consequent impacts on marine ecosystems (Kroeker et al., 2013). Climate change models predict a global increase in mean sea surface temperature of 1–4°C and a decrease of ~0.3–0.4 pH by the year 2100 (IPCC, 2014, 2019). The Southern Ocean, the water masses extending south of the Polar Front to the coasts of the Antarctic continent, is predicted to be among the first and most highly affected regions by ocean acidification due to increased solubility of CO_2_ at low temperatures leading to low carbonate ion concentrations ([CO_3_ ^2−^]) (Fabry et al., 2009; Orr et al., 2005). Ocean acidification is also shallowing the saturation horizon, which is the depth below which calcium carbonate (CaCO_3_) minerals (such as aragonite and calcite) can dissolve. It is anticipated that surface waters in some Southern Ocean regions will become undersaturated with respect to aragonite and calcite by 2050 and 2095, respectively (McNeil & Matear, 2008; Orr et al., 2005), and that 70% of the Southern Ocean could be aragonite undersaturated by 2100 (Hauri et al., 2016). Due to the low interannual variability in surface ocean [CO_3_ ^2−^] of this region (Conrad & Lovenduski, 2015), many organisms may be unable to adapt rapidly to these changing conditions. Some parts of the continent, such as the Western Antarctic Peninsula, are also among the fastest warming regions globally (Steig et al., 2009). The Southern Ocean acts as a sentinel region for evaluating the impacts of ocean warming and acidification on marine organisms and for making predictions about future impacts at lower latitudes where ocean acidification is expected to occur later (Andersson et al., 2008; Fabry et al., 2009; Figuerola et al., 2021).

Understanding how marine organisms can cope with these environmental changes is essential for predicting the future of marine ecosystems. Marine calcifiers, organisms which build their skeletons from CaCO_3_, may be particularly sensitive to these threats. Increased *p*CO_2_ and decreased pH, caused by ocean acidification, increase the solubility of CaCO_3_ skeletons (Andersson et al., 2008; Fabry, 2008; Figuerola et al., 2021). The ability of marine calcifiers to respond to these rapid environmental stresses may, therefore, be compromised, especially for Southern Ocean taxa, which are often characterized by having lower metabolic, development and growth rates than related species from warmer latitudes (Peck, 2018; Peck, 2001, 2005). Southern Ocean taxa that deposit more soluble CaCO_3_ mineral phases such as aragonite and high Mg-calcite could be at greater risk to changes in ocean carbonate chemistry than species secreting a mixture of calcite and aragonite (bimineralic) or lower Mg-calcite levels (Andersson et al., 2008; Figuerola et al., 2021; Lebrato et al., 2013; McClintock et al., 2011, 2017). For instance, common habitat-forming or key predator taxa, which have important roles in Antarctic ecosystems, have aragonitic (e.g. scleractinian cold-water corals) or high Mg-calcite skeletons (e.g. echinoderms and bryozoans) (Bax & Cairns, 2014; Figuerola et al., 2019; McClintock et al., 2011; Turley et al., 2007). The negative ocean acidification impact on these taxa could thus lead to the loss of habitat complexity and resilience in these ecosystems.

Ocean warming may even accelerate skeletal solubility as a positive correlation between Mg-calcite and/or aragonite and seawater temperature has been found in diverse phyla such as molluscs, serpulid annelids, foraminifera, corals, echinoderms, bryozoans and crustaceans (Chang et al., 2004; Chave, 1954; Cohen & Branch, 1992; Lowenstam, 1954). Seawater temperature seems to affect the spatial distribution of marine calcifiers, with higher abundances of species depositing aragonite and/or higher Mg-calcite levels in the tropics, although this pattern has yet to be rigorously tested in some groups such as bryozoans.

In addition, there remain uncertainties in the predicted effects of ocean warming on marine calcifiers because diverse environmental factors (e.g. salinity, pH, CaCO_3_ saturation state, seawater Mg/Ca ratio and diet) and biological (e.g. skeletal growth rate) may interfere in the calcification process, especially on a local scale (Amsler et al., 2010; Borremans et al., 2009; Duquette, Vohra, et al., 2018; Figuerola et al., 2015; Krzeminska et al., 2016; Kołbuk et al. 2019; Krzemińska et al., 2022; Lebrato et al., 2016; Lowenstam & Weiner, 1989; Loxton, Kuklinski, Barnes, et al., 2014; Ries, 2010). For instance, variability in seawater Mg/Ca ratios (Lebrato et al., 2020) is known to affect the skeletal Mg-calcite and mineralogy of a range of species (Ries, 2010; Kołbuk et al. 2021). Among biological processes, “vital effects” may distort or mask any signal of environmental effects on mineralogy (Weiner & Dove, 2003). Plasticity is also a mechanism by which calcifiers such as molluscs may adapt to environmental changes, as observed when organisms are exposed to different experimental conditions (Figuerola et al., 2021; Hancock et al., 2020). Therefore, more studies are needed on a wider range of phyla and species with broad geographical ranges, skeletal mineralogies, biological traits and physiological processes.

Among calcifying taxa, bryozoans are abundant and speciose suspension-feeding aquatic invertebrates worldwide (>6000 extant species; Bock and Gordon, 2013), being present from the intertidal to the deep sea (e.g. Barnes and Kuklinski, 2010; Figuerola et al., 2018). They are dominant members of many benthic communities, including the Antarctic region (Barnes & Kuklinski, 2010; Figuerola et al., 2012). Many species are major carbonate sediment producers, thus being well represented in the fossil record (Taylor, 2020) and good model systems for understanding ecological and evolutionary processes (Orr et al., 2022). The calcareous skeletons of bryozoans can be entirely calcitic, aragonitic or bimineralic, and incorporate varying amounts of Mg in their calcite, ranging from low Mg-calcite (LMC; <4 wt% MgCO_3_), through intermediate Mg-calcite (IMC; 4−8%), to high Mg-calcite (HMC; >8%) following the categories proposed by Rucker and Carver (1969) (see Taylor et al., 2015).

Early studies exploring marine invertebrate mineralogy proved cheilostome mineralogy to be highly variable, unlike other taxa, thus having great potential for reconstruction changes (Lowenstam, 1954; Poluzzi & Sartori, 1975; Rucker & Carver, 1969). The mineralogical studies have persisted and proliferated even to today (e.g., Bone and James, 1993; Borisenko and Gontar, 1991; Smith et al., 1998). In recent years significant efforts have been put into increasing data on skeletal mineralogy in a wide range of bryozoan species from Arctic (Iglikowska et al., 2020; Krzemińska et al., 2022; Kuklinski & Taylor, 2009; Piwoni-Piórewicz et al., 2019), temperate (e.g., Loxton et al., 2014b; Smith and Clark, 2010; Steger and Smith, 2005), tropical (Taylor et al., 2016; Taylor & Di Martino, 2014), and Antarctic (Figuerola et al., 2015, 2019; Krzeminska et al., 2016; Loxton et al., 2013; Loxton, Kuklinski, Barnes, et al., 2014) regions. However, several reviews have pointed out that mineralogy is known only for a small proportion of living species at all latitudes (Smith et al., 2006; Taylor et al., 2009).

Southern Ocean bryozoans, which comprise over 300 species, exhibit interspecific variations in the Mg-calcite of their skeletons, and wide geographic and depth distributions (Barnes & Kuklinski, 2010; De Broyer et al., 2011; Figuerola et al., 2012, 2017; Hayward, 1995; Krzeminska et al., 2016), making them suitable indicators to monitor the effects of ocean warming and acidification in a hotspot for global change. As ecosystem engineers, this group also creates habitat heterogeneity and refugia for many associated macro- and micro-invertebrates and thus are likely to boost ecological resilience (Santagata et al., 2018). In this study, we present the largest carbonate mineralogical dataset for Southern Ocean bryozoans to date. To improve our understanding of how bryozoans might respond to ocean warming and acidification globally, we then employ rigorous correlative statistical analyses to assess latitudinal and seafloor temperature patterns of skeletal mineralogy. We hypothesized that seawater temperature is the main driver on influencing mineralogical composition in bryozoans at a global scale as maintaining aragonitic and HMC skeletons is more energetically demanding in cold waters. Thus, we expect that the seawater temperature affects spatial distribution of bryozoans. To our knowledge, this is the first study to test this hypothesis using a seafloor temperature data set and species occurrences combined with new and published mineralogical data.

## 2 MATERIAL AND METHODS

### 2.1 Study area

Bryozoan specimens were collected from East Antarctica using beam trawl during the CEAMARC cruise on the RSV ‘Aurora Australis’ (2007 to 2009). The study area covers part of the region Terre Adélie (141−142° E) and George V Land (142−145° E). Collection depths ranged from 138−662 m. Sampling sites were geo-referenced using GPS and depth was recorded at each site (Fig. 1; Table S1; Tables S1–4 are given in the Dryad Digital Repository: https://datadryad.org/stash/share/3ZheuFuQESDCw-AQUt10tE7f2ol5uIDSe0Vv5laUtBQ). On the research vessel, all bryozoan samples were preserved in 95% ethanol. Bryozoan colonies were identified to the lowest possible taxonomic rank in the laboratory using an optical microscope and a taxonomic reference guide (Hayward, 1995). To confirm identification, a subset of specimens was bleached for 1−4 hours in a 10% solution of sodium hypochlorite (domestic bleach, diluted using tap water), rinsed in freshwater, air-dried, and examined using a LEO SEM at the Natural History Museum in London (Fig. 2).

**Fig. 1.**
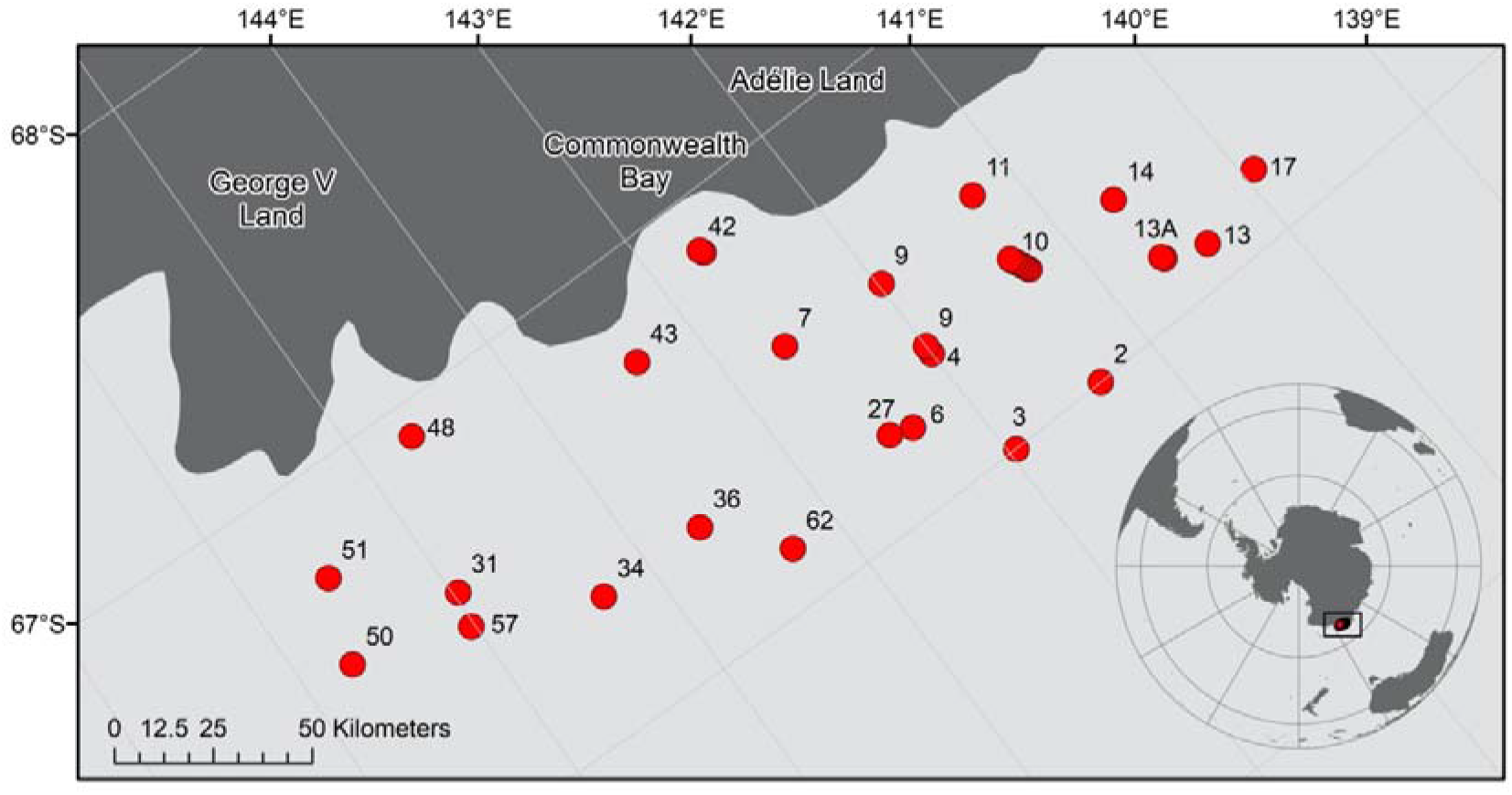
Map of the study area (Terre Adélie and George V Land) in East Antarctica and the sampling stations.

**Fig. 2.**
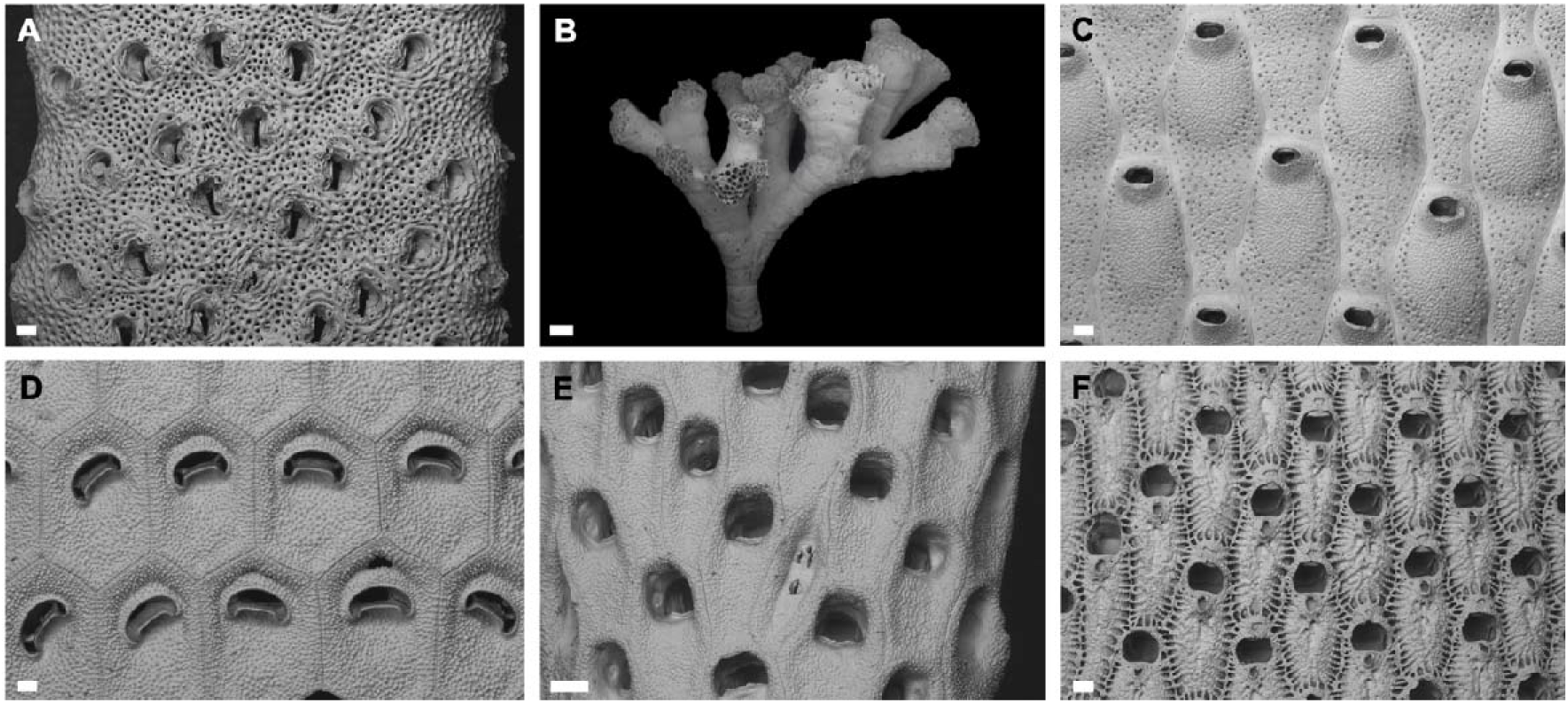
SEM images of 6 selected bryozoan species studied from East Antarctica. (A) *Systenopora contracta*, zooids on branch surface (scale bar = 200 μm); (B) *Fasciculipora ramosa*, colony (scale bar = 1 mm); (C) *Lageneschara lyrulata*, group of zooids (scale bar = 200 μm); (D) *Melicerita obliqua*, zooids from near centre of branch (scale bar = 100 μm); (E) *Chondriovelum adeliense*, group of zooids (scale bar = 200 μm); (F) *Astochoporella cassidula*, group of zooids (scale bar = 200 μm).

### 2.2 Mineralogical analyses

Specimens were cleaned carefully of epibionts using a scalpel blade to avoid mineralogical contamination. If possible a piece was cut from the growing edge of each specimen and air dried as described in previous studies (Figuerola et al., 2015; Kuklinski & Taylor, 2009). Otherwise any available part of colony was used for analysis. The pieces were powdered using an agate pestle and mortar. Mineralogical analyses were carried out at the Natural History Museum using a high-precision Enraf-Nonius X-ray diffractometer (XRD) equipped with an INEL CPS-120 Curved Position Sensitive detector (−120°, 2-theta) and a cobalt X-ray source. Operating conditions of the cobalt source were 40 kV and 40 mA. The tilt angle between source and sample was 5.9°. The horizontal slit system was set to 0.14 mm to confine the X-ray beam to pure cobalt Kα1. An internal standard (NaCl) was added to the samples to correct for sample displacement. The samples were rotated during the measurements to improve the randomness of grain orientations in the X-ray beam. The 2-theta linearity of the detector was calibrated using Y_2_O_3_ (yttrium (III) oxide) (99.9% BDH Laboratory Chemicals) and SRM 640 silicon powder (NIST) and the calibration curve was fitted using a least-squares cubic spline function. Peak positions were calibrated on Y_2_O_3_ data from Mitric et al. (1997). Bryozoan skeleton was compared with external standard of calcite (Iceland spar) by fitting peak intensities of the sample patterns and the standard patterns. The wt% MgCO_3_ in calcite was calculated by measuring the position of the d_104_ peak, assuming a linear interpolation between CaCO_3_ and MgCO_3_ (Mackenzie et al., 1983). A linear trend of d_104_ vs. mol% MgCO_3_ can be observed in the range between 0 and 17 mol% MgCO_3_ (Mackenzie et al. 1983); all analyses from the present study fall within this range. This composition information is accurate to within 2% on a well-calibrated instrument (Kuklinski & Taylor, 2009).

### 2.3 Temperature data

The seafloor temperature data set employed was that presented in Clarke et al. (2009). Seawater temperature values were taken from the World Ocean Atlas 2005, a data product of the Ocean Climate Laboratory of the U.S. National Oceanographic Data Centre (http://www.nodc.noaa.gov/OC5/WOA05/pr_woa05.html). Global mean annual ocean *in situ* temperature data interpolated into 33 standard vertical intervals (depth ranges) at a 1° spatial resolution were overlain onto a bathymetry grid (ETOPO1) (Amante & Eakins, 2009). Seawater temperature data were then extracted when the depth range coincided with the bathymetry of the seafloor to create a seawater temperature data set. Potential temperature was calculated according to the UNESCO equation of state for seawater and the International Temperature Scale of 1990 (ITS-90) specified by the International Committee of Weights and Measures. A nearest neighbor interpolation was then used to create a seafloor temperature grid for both *in situ* and potential temperature in ArcGIS. Temperatures were extrapolated for each bryozoan occurrence record (latitude and longitude) to assign a temperature value.

### 2.4 Literature search and bryozoan species distributions

We integrated new data from the current study with recent (e.g., Krzeminska et al., 2016; Loxton et al., 2018) and older mineralogy data from cheilostome and cyclostome bryozoans worldwide previously summarized by Smith et al. (2006) and Taylor et al. (2009). Available mineralogical data from genera with undetermined species, and also known species from the White Sea which has unusual conditions that may mask temperature influence (Krzemińska et al., 2022), were removed from the analyses (see Tables S2–S4). The global dataset comprised 549 species (102 Southern Ocean species, excluding six genera with indeterminate species). Calcite composition was classified as LMC, IMC and HMC. Data previously reported as mol% MgCO_3_ were converted to weight% (wt%) MgCO_3_ to facilitate comparisons.

To generate the mineral type and skeletal Mg-calcite distribution maps and graphs, we downloaded occurrence data for all living bryozoan species from the Global Biodiversity Information Facility (GBIF, n.d.) (GBIF.org (28 January 2022) GBIF Occurrence Download https://doi.org/10.15468/dl.59nzhf). We used our list of 549 species with available mineralogical data (with species names verified using the World Register of Marine Species Taxon Match tool-https://www.marinespecies.org/aphia.php?p=match) to filter the downloaded GBIF data. Duplicate occurrence records (repeated combinations of latitude, longitude, and scientific name) and any occurrences on land were removed. The filtered GBIF dataset comprised 96,241 occurrences for the 493 selected species that had occurrence records within GBIF (Table S5).

### 2.5 Statistical analysis

Coefficients of determination (R^2^) for the regressions and *p* values were calculated to evaluate correlative relationships between latitude and temperature and mineralogical composition. Data were analysed and figures constructed in: R v. 4.1.0/Rstudio v. 1.4.1717, using the R packages Tidyverse (ggplot2) and Ggally v. 2.1.2 (Schloerke et al., 2021; Wickham et al., 2019). Statistical significance was set at *p* < 0.05.

## 3 RESULTS

### 3.1 Skeletal mineralogy of the Southern Ocean and warmer regions

Among the new samples (n = 126) analysed in the present study, 37 Antarctic bryozoan species had skeletons consisting of calcite and were characterized by a wide range of Mg-calcite in the skeleton (1.52−7.55 wt% MgCO_3_; Fig. 3; Table S2). The mean Mg-calcite levels for all analysed samples was 4.15 ± 1.28 wt% MgCO_3_. Fifty-seven percent of the species we examined were categorized as IMC (21 species), whereas the remaining 43% were categorized as LMC (16 species; Fig. 3).

**Fig. 3.**
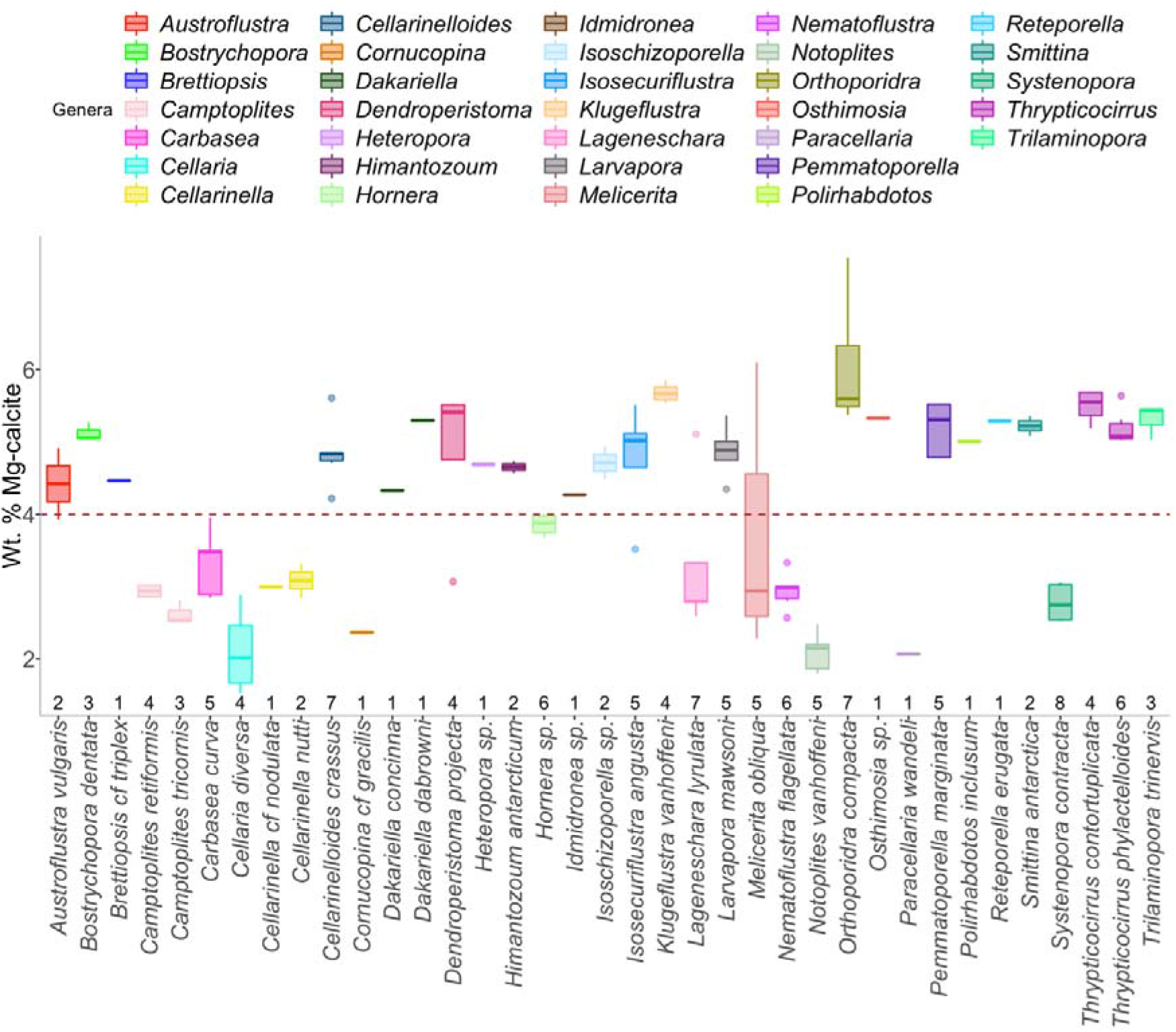
Mean values (±SD) of wt% MgCO_3_ in skeletal calcite of the new species analysed. Boxes show standard deviation around mean (mid-line), tail indicates range, and dots are outliers. Data for species are presented alphabetically left to right. The horizontal dashed line indicates the upper boundary of the LMC. Numbers are sample sizes.

The resulting data set, when combined similar data from the literature, included 102 Southern Ocean species (n = 1415; Table S3). The average Mg-calcite from all analysed samples was 4.54 ± 1.32 wt% MgCO_3_. Most species fell into the IMC category (67%; 68 species) while the remaining were in the LMC category (32%; 33 species) apart from *Beania erecta* Waters, 1904 which was in the HMC category (1%; 8.03 ± 1.56 wt% MgCO_3_) (Fig. 4A; Table S3).

**Fig. 4.**
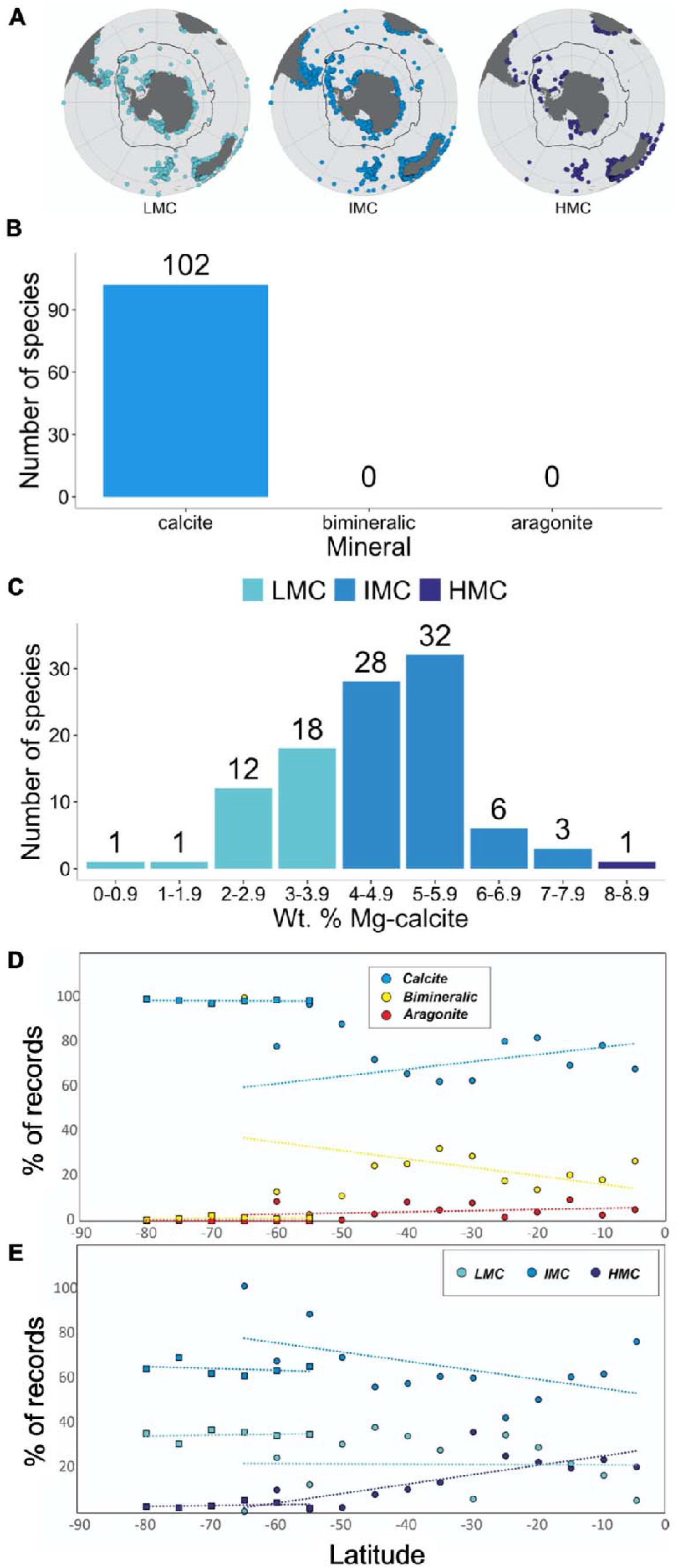
Distribution patterns of the mineral type and of the skeletal Mg-calcite of bryozoans in the Southern Hemisphere. (A) Maps of the Southern Ocean and neighbouring regions showing the Polar Front (PF; black line) and the distribution patterns of Mg-calcite in bryozoan skeletons; Frequency distributions for mineral type (B) and (C) skeletal Mg-calcite in 1415 specimens belonging to 102 species from the Southern Ocean; Relationships between latitude (D) and (E) mineralogical composition in the Southern Hemisphere. LMC: low Mg-calcite; IMC: intermediate Mg-calcite; HMC: high Mg-calcite. South of PF (squares): Calcite = R^2^ = 0.0166, *p* = 0.81, y = 98 − 0.009X; Bimineralic = R^2^ = 0.0334, *p* = 0.73, y = 2.2 + 0.013X; Aragonite = R^2^ = 0.1543, *p* = 0.44, y = −0.24 − 0.0041X. North of PF (circles): Calcite = R^2^ = 0.0743, *p* = 0.37, y = 81 + 0.33X; Bimineralic = R^2^ = 0.0957, *p* = 0.3, y = 13 −0.38X; Aragonite = R^2^ = 0.0831, *p* = 0.34, y = 6.1 + 0.051X. South of PF (squares): LMC = R^2^ = 0.0387, *p* = 0.71, y = 37 + 0.045X; IMC = R^2^ = 0.0698, *p* = 0.62, y = 58 − 0.081X; HMC = R^2^ = 0.0634, *p* = 0.63, y = 5.2 + 0.036X. North of PF (circles): LMC = R^2^ = 0.0004, *p* = 0.95, y = 21 − 0.012X; IMC = R^2^ = 0.2571, *p* = 0.077, y = 50 − 0.41X; HMC = R^2^ = 0.5802, *p* = 0.0025, y = 29 + 0.42X.

Globally, 72% (397 species) of a total of 549 species were calcitic, 22% (122 species) bimineralic and 6% (30 species) aragonitic (Fig. 5A). Most bryozoan species had IMC (65%; 322 species) while the remaining had LMC (27%; 133 species) or HMC (8%; 41 species) (Fig.6A).

**Fig. 5.**
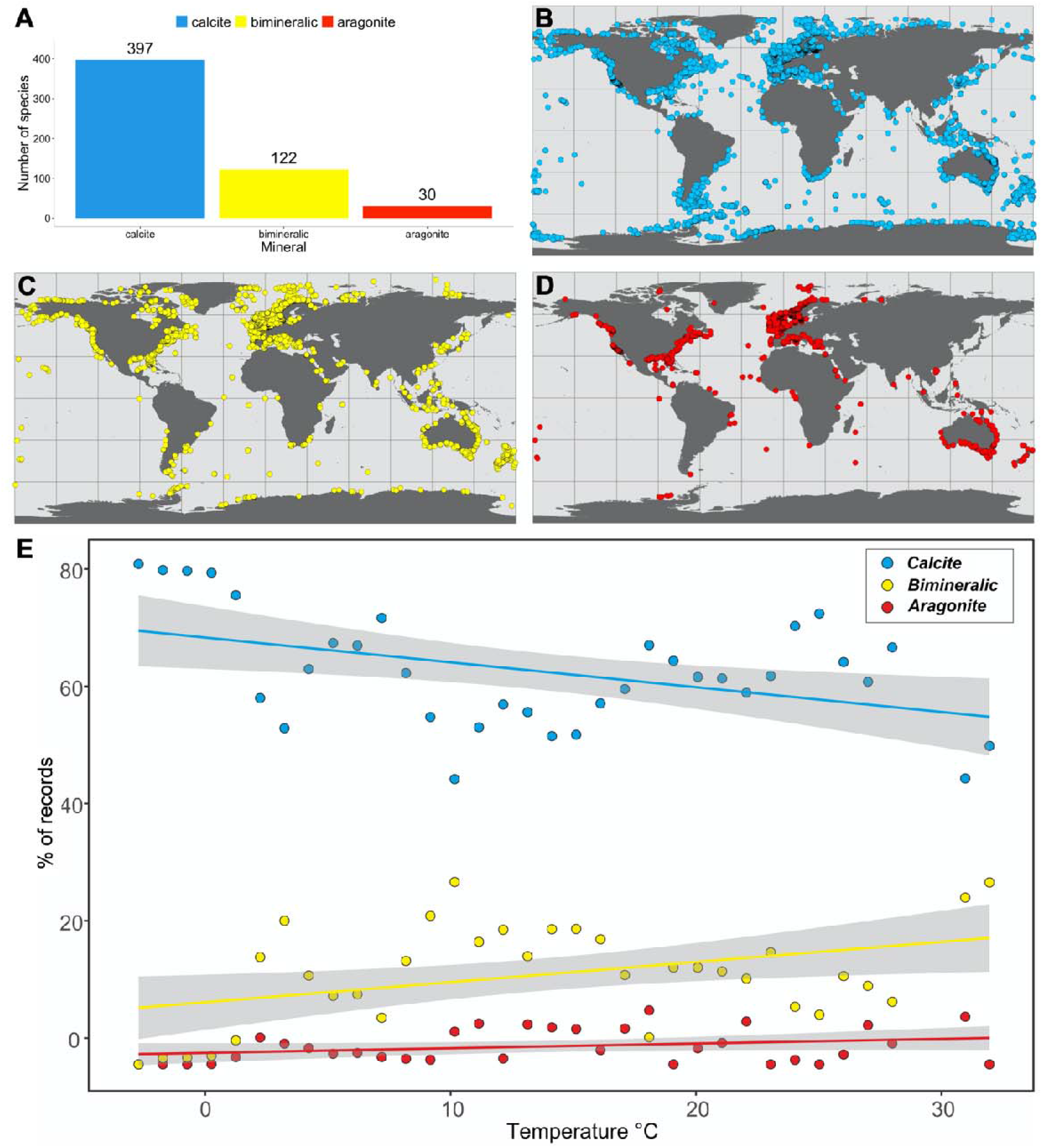
Global distribution patterns of the mineral type of bryozoans. (A) Global frequency distributions for mineral type; (B) Global distribution patterns of calcite (blue), (C) bimineralic (yellow) and (D) aragonite (red); E) Relationships between seawater temperature and mineral type in the Southern Hemisphere. Shading bands represent standard errors on linear regressions. Calcite = R^2^ = 0.1882, *p* = 0.01, y = 85 − 0.49X; Bimineralic = R^2^ = 0.1648, *p* = 0.017, y = 13 + 0.4X; Aragonite = R^2^ = 0.079, *p* = 0.11, y = 2.3 + 0.092X.

### 3.2 Global-scale latitudinal patterns of skeletal mineralogy

The proportions of occurrence records of different mineral types remained stable south of the Polar Front, with aragonite almost completely absent, bimineralic occurrence records less than 3% with most occurrence records (97%) dominated by calcite. North of the Polar Front was also dominated by calcite but to a lower level (~62–97%), higher levels of bimineralic skeletons (~3–32% and 100% for a single occurrence record south of 60° S) (Table S5). There was also a trend of increasing aragonite (from zero up to a maximum of ~9.5%) with decreasing latitude (Fig. 4C). There was a significant increase in the percentage of bimineralic occurrence records (R^2^ = 0.16; *p* = 0.017) and decrease of calcitic occurrence records (R^2^ = 0.19; *p* = 0.01) with an increase in seafloor water temperature (Fig. 5E). Aragonitic occurrence records were absent below 0°C and very low below 1°C (0.05%), with no significant trend in waters above that temperature (*p* = 0.11) with proportions ranging between 1% and 11% of the total number of occurrences.

The proportions of LMC, IMC and HMC occurrence records did not vary south of the Polar Front either (Fig. 4C). IMC dominated occurrence records (60–68%), 30% to 36% of occurrence records were LMC and HMC was only present in very low proportions (1.5–5%). However, HMC species significantly increased in prevalence from the Polar Front towards lower latitudes (R^2^ = 0.58; *p* = 0.0025; Fig. 4C) and with increasing seawater temperatures (R^2^ = 0.48; *p* = 3.6e^−06^; Fig. 6E). IMC species had the opposite trend, with a decrease towards lower latitudes (R^2^ = 0.26; *p* = 0.077; Fig. 4C) and a significant decrease with increasing seawater temperatures (R^2^ = 0.45; *p* = 1.1e^−06^; Fig. 6E). No significant trend was found in LMC species north of the Polar Front with latitude or with temperature.

**Fig. 6.**
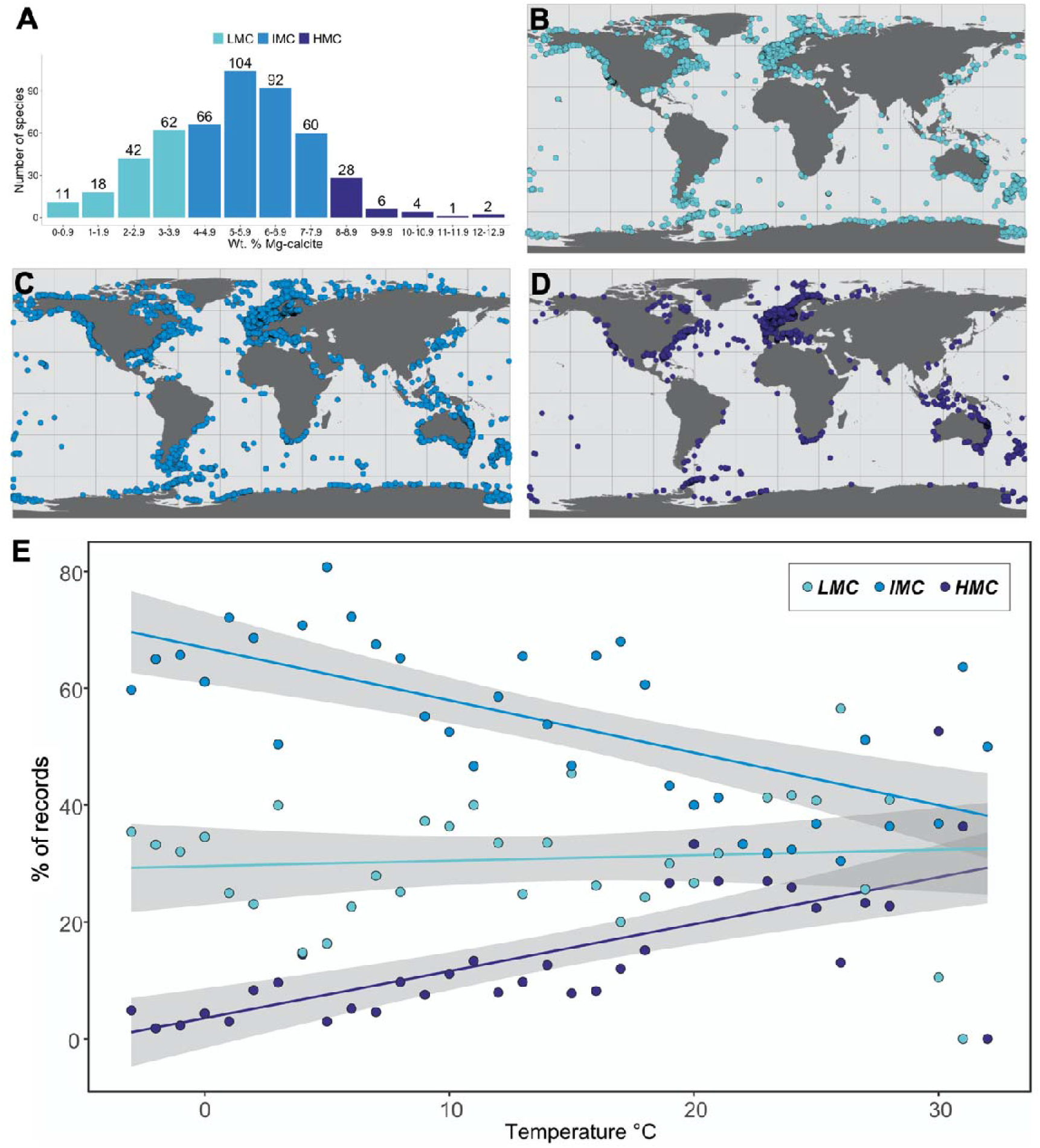
Global distribution patterns of the skeletal Mg-calcite of bryozoans. (A) Global frequency distributions for skeletal Mg-calcite; (B) Global distribution patterns of low Mg-calcite (LMC; blue), (C) intermediate Mg-calcite (IMC; yellow) and (D) high Mg-calcite (HMC; red); (E) Relationships between seawater temperature and mineralogical composition in the Southern Hemisphere. Shading bands represent standard errors on linear regressions. LMC = R^2^ = 0.0067, *p* = 0.62, y = 30 + 0.094X; IMC = R^2^ = 0.4488, *p* = 1.1e-05, y = 67 − 0.9X; HMC = R^2^ = 0.4828, *p* = 3.6e-06, y = 3.5 + 0.8X.

## 4. DISCUSSION

### 4.1 Seawater temperature as a likely driver influencing mineralogical composition at a global scale

In the present study we have compiled, and in some cases analysed, the largest carbonate mineralogical and distribution dataset for Southern Ocean bryozoans to date, including data of approximately one-third of the known extant bryozoan species from Antarctica based on those listed in the Register of Antarctic Marine Species database (102 of 335 species) (De Broyer et al., 2022). This dataset thus increases our knowledge of the potential effects of global change on Antarctic bryozoans, an important phylum of invertebrates in the Antarctic benthos which creates framework habitats that enhance biodiversity and produces large quantities of carbonate sediments, thus making up significant blue carbon sinks (Barnes, 2018). These data provide a useful baseline for understanding the vulnerability of different bryozoan species and make predictions of the effects of ocean acidification on benthic habitats and sediments.

Our results confirm that all Southern Ocean species have monomineralic calcite skeletons, as previously revealed by several studies on bryozoan skeletal mineralogy (Figs. 3 and 4) (Borisenko & Gontar, 1991; Figuerola et al., 2015, 2019; Krzeminska et al., 2016; Loxton et al., 2013; Loxton, Kuklinski, Barnes, et al., 2014; Taylor et al., 2009). While occasional analyses of specimens belonging to the bryozoan species *Chondriovelum* [*Labioporella*] *adeliensis* (Livingstone, 1928) and *Isosecuriflustra angusta* (Kluge, 1914) detected aragonite (Borisenko & Gontar, 1991; Loxton et al., 2013), recent analyses including the current study contradict these findings and provide evidence of their monomineralic calcitic nature (Krzeminska et al., 2016; Taylor et al., 2009). We suggest that these detections of aragonite are likely due to sample contamination by epibionts or sediment (Taylor et al., 2015).

Our data illustrate that the skeletons of most Southern Ocean species of bryozoans are formed of IMC, some of LMC and a few of HMC, consistent with previous global data (Figs. 3 and 4; Table S3) (Smith et al., 2006; Taylor et al., 2009). However, we found higher IMC (67 vs 65*−*58%) and LMC (32 vs 29*−*27%) percentages and a lower HMC (1 vs 13*−*8%) percentage compared to those from our own and previous global data this was as expected due to the higher solubility of HMC in cold waters (Fig. 6A) (Davis et al., 2000; Mucci, 1983; Smith et al., 2006; Taylor et al., 2009).

Our findings also reveal that the proportions of skeletal aragonite, bimineralic mixtures and calcite remained stable south of the Polar Front consistent with the low and nearly constant temperatures found in the Southern Ocean (Fig. 4 C) (Moore et al., 1999). From the Polar Front boundary towards lower latitudes, we did not detect any clear general latitudinal trend (Fig. 4 C) as previously found in studies that did not extrapolate values onto wider occurrence data (e.g. Taylor et al., 2009). However, when we plotted our mineralogical findings against the seafloor temperature, there were an increase in the proportions of bimineralic occurrence records (R^2^ = 0.16; *p* = 0.017) and a decrease in the proportion of calcitic occurrence records (R^2^ = 0.19; *p* = 0.01) with increasing temperature consistent with decreased costs of skeletal production (Fig. 5E). The lack of a significant increase in aragonite occurrence records with increasing temperature (R^2^ = 0.079; *p* = 0.11) is potentially due to a lack of published mineralogical studies of recent bryozoans from the tropics, where most species with aragonitic skeletons occur (Taylor et al., 2009, 2016). However, species with aragonitic skeletons are absent from seas with temperatures of zero degrees centigrade or below, indicating that this is some sort of temperature threshold or limit for aragonitic species.

In addition, two cosmopolitan aragonitic species, *Membranipora membranacea* (Linnaeus, 1767), which had high occurrence records (6759), and *Hippothoa divaricata* Lamouroux, 1821, which had some Antarctic occurrence records (n = 21), could mask the general trend (Table S5). These species could also switch in mineralogy in higher latitudes, although they were not found below zero degrees Celsius. Some mineralogical analyses of *M. membranacea* recovered calcite (Taylor et al., 2009), although it is not clear if it is due to taxonomic misidentification.

Interestingly, the only Antarctic occurrence records of *H. divaricata* are from the Western Antarctic Peninsula, where the water is warmer than other Antarctic regions. We thus suggest that zero degrees Celsius could be a better cut off point, above which aragonite does not show a significant trend towards the equator. However, further taxonomic studies are needed to confirm the identification of this species. It should be also taken into account that our analyses make the assumption that species do not switch mineralogy, although it has been found in some bryozoan species (e.g. Smith and Girvan, 2010), and might impact some of our findings.

Mg-calcite remained low and constant south of the Polar Front and significantly increased towards lower latitudes (HMC R^2^ = 0.58; *p* = 0.003) and with increasing seafloor water temperatures (HMC R^2^ = 0.48; *p* < 0.001), consistent with our hypothesis that seawater temperature influence on bimineralization seems to affect the spatial distribution of bryozoans. For the first time we thus provide evidence that seafloor water temperature is likely a significant driver influencing mineralogical composition in bryozoans at a global scale, with an increase of HMC and bimineralic species in warmer waters. Previous studies, lacking an assessment of temperature, revealed an inverse correlation between Mg-calcite and latitude in different calcifying marine invertebrates (Andersson et al., 2008; Duquette, Halanych, et al., 2018; McClintock et al., 2011). For instance, this general pattern was observed in echinoderm species with LMC skeletons only found in Antarctica although most species have HMC skeletons (Duquette, Halanych, et al., 2018; Figuerola et al., 2021). Smith et al. (2016) also found a similar trend in different life stages and nine skeletal elements of echinoids. Other taxa also have LMC shells at higher latitudes such as Antarctic brachiopods (Barnes & Peck, 1996). In the case of bryozoans, a latitudinal gradient in cheilostome mineralogy, in which aragonite and Mg-calcite increase towards the tropics, was also observed (Krzeminska et al., 2016; Piwoni-Piórewicz et al., 2019; Taylor et al., 2009). Kuklinski & Taylor (2008, 2009) also compared congeneric species across a latitudinal transect and corroborated a shift towards aragonitic and bimineralic skeletons from the Arctic into lower latitudes. Our assessment of latitudinal patterns also provide support for the viability of using latitudinal gradients to infer bryozoan species’ responses to seawater temperature although more tropical species should be analysed mineralogically to confirm this hypothesis.

Our data may reflect several adaptive advantages of secreting monomineralic calcite and less soluble CaCO_3_ mineral phases (e.g. LMC skeletons) in cold waters bryozoans such as more rapid secretion of large zooids/colonies, more energetically efficient skeleton secretion and reduction in skeleton dissolution as previously suggested for Arctic bryozoans (Kuklinski & Taylor, 2009). Consistent with the high energetic costs to maintain calcified skeletons, especially those with HMC, in cold waters, a decrease of skeletal investment with latitude and decreasing temperature has been also noted in several taxa such as gastropods and echinoids (Watson et al., 2017). Globally, an increase of bryozoans depositing stable carbonate minerals such as calcite and LMC is thus expected as previously suggested by Andersson et al. (2008).

The energetic cost of calcification due to the increasing solubility of CaCO_3_ with decreasing temperature may also lead to compensatory mechanisms in diverse species, likely improving their resilience to global change. For instance, a recent study showed mussels produce thinner shells with a higher organic content in the polar region compared to temperate latitudes (Telesca et al., 2019). Some brachiopod and coral species can also have the ability to form thicker shells and denser skeletons, respectively, under acidified conditions (Cross et al., 2019; Teixidó et al., 2020). However, these changes in the calcification may not be sufficient to overcome impacts of both warming and acidification.

At a local scale, it remains unclear what the effect of seawater temperature and other environmental (e.g. salinity, pH and the seawater Mg/Ca ratio) or biotic factors have on the skeleton of bryozoans. For instance, the expected positive trends of increasing aragonite or Mg-calcite with seawater temperature were not found in some polar and temperate bryozoan species although the authors argued that the existing variability in seawater temperature might be too low in these instances to significantly affect skeletal Mg-calcite (Figuerola et al., 2015; Iglikowska et al., 2020; Loxton, Kuklinski, Barnes, et al., 2014; Loxton, Kuklinski, Najorka, et al., 2014). In addition, local intraspecific and/or interspecific variation was previously found in a range of bryozoan species, which were not related with temperature changes (Figuerola et al., 2015, 2019; Loxton, Kuklinski, Barnes, et al., 2014; Loxton, Kuklinski, Najorka, et al., 2014) suggesting other biological and/or local environmental factors remain important. This could be the case of some Antarctic and temperate bryozoan species, which exhibited a decrease in skeletal Mg-calcite at lower salinities (Figuerola et al., 2015; Loxton, Kuklinski, Najorka, et al., 2014). In contrast, Loxton et al. (2014a) did not found any relationship between the skeletal Mg-calcite and salinity in four Antarctic bryozoans. Several studies investigating the effect of temperature and salinity on Mg/Ca ratios in other calcifying groups also revealed interspecific variation. While a positive correlation between salinity and skeletal Mg/Ca ratio was found in some foraminifer and echinoderm species – although Mg/Ca responses to changes in temperature are more strongly (Borremans et al., 2009; Dissard et al., 2010; Hönisch et al., 2013), no correlation with salinity was showed in other foraminifer and echinoderm species (Hermans et al., 2010; Geerken et al., 2018). Further laboratory and field research should thus attempt to tease apart the influence of seawater temperature and other factors such as salinity on bryozoan mineralogy.

### 4.2 Southern Ocean bellwethers for global change impacts on marine calcifiers

Southern Ocean bryozoans are likely to be particularly vulnerable to global change impacts as most cyclostomes and cheilostomes release lecithotrophic short-lived larvae (Jackson, 1986) and may thus not be able to shift their distribution to more favourable habitats. Our findings suggest that the combined effects of ocean warming and acidification may be seen first in HMC species (e.g. *Beania erecta*) inhabiting in Antarctic coastal areas that have rapidly warmed in recent years such as the Western Antarctic Peninsula.

Bryozoan species secreting IMC and even LMC skeletons may also be at increased risk to global change in the near-future, as their skeletons could show an increase in Mg-calcite with warmer seawater temperatures (Figuerola et al., 2019). Bryozoan species of the genus *Pentapora* showed an increase of Mg-calcite in parts of the skeleton formed during the summer (Lombardi et al., 2008). Colonies of two common Mediterranean bryozoans, *Myriapora truncata* (Pallas, 1766) and *Pentapora fascialis* (Pallas, 1766), also exhibited higher Mg-calcite values when exposed to experimental high temperatures compared to those of control conditions. These differences only approached significance perhaps due to the short time period of the controlled temperature exposure period (Pagès-Escolà et al., 2018). Skeleton dissolution was also observed in the bryozoan *Jellyella tuberculata* (Bosc, 1802), when colonies were exposed to a combination of reduced pH and elevated temperature, likely due to an increase in levels of skeletal Mg-calcite with increased temperature, making skeletons more susceptible to dissolution at low pH (Swezey et al., 2017). However, these findings contrast with those of Duquette et al. (2018b) who found that a chronic exposure to elevated temperature resulted in a significant reduction in the Mg-calcite of the sea urchin *Lytechinus variegatus* (Lamarck, 1816). Clearly, in a big picture, more experimental and *in-situ* studies that combine near-future predicted elevated temperature and reduced pH (Cummings et al., 2019; Stark et al., 2018) are needed to predict the bryozoans that will be “winners” and “losers” a determination that will be particularly important for those species that are key ecosystem engineers, whose loss could lead to a decline of resilience in benthic Antarctic ecosystems.

## ACKNOWLEDGMENTS

We would like to acknowledge the voyage leader, Martin Riddle, the crew and the captain of the RV Aurora Australis. The CAML-CEAMARC cruise of RV Aurora Australis (IPY project n°53) were supported by the Australian Antarctic Division, the Japanese Science Foundation, the French polar institute IPEV and the Muséum National d’Histoire Naturelle). We thank Dr Steve J. Roberts for helping with the figures 6 and 7, Jen Najorka for help with mineralogical analysis and Dr Paul D. Taylor and three anonymous reviewers for their valuable comments and edits. The study was supported by National Science Centre Poland (project contracts 2016/23/B/ST10/01936 and 2021/40/C/ST10/00225). BF has received funding from the postdoctoral fellowships programme *Beatriu de Pinós* funded by the Secretary of Universities and Research (Government of Catalonia) and by the Horizon 2020 programme of research and innovation of the European Union under the Marie Skłodowska-Curie grant agreement No 801370 (Incorporation grant 2019 BP 00183). This work has been developed in the framework of MedCalRes project Grant PID2021-125323OA-I00 funded by MCIN/AEI/ 10.13039/501100011033 and by “ERDF A way of making Europe”. With the institutional support of the ‘Severo Ochoa Centre of Excellence’ accreditation (CEX2019-000928-S).

